# Development of a chemiluminescence immunoassay to accurately detect African swine fever virus antibodies in serum

**DOI:** 10.1101/2021.02.28.433302

**Authors:** Yong Yang, Changjie Lv, Junqing Fan, Ya Zhao, Lili Jiang, Xiaomei Sun, Qiang Zhang, Meilin Jin

## Abstract

The outbreak of African swine fever (ASF) has caused significant economic losses to animal husbandry worldwide. Currently, there is no effective vaccine or treatment available to control the disease, and therefore, efficient disease control is dependent on early detection and diagnosis of ASF virus (ASFV). In this study, a chemiluminescent immunoassay (CLIA) was developed using the ASFV protein p54 as a serum diagnostic antigen and an anti-p54 monoclonal antibody. After optimizing the working parameters of the CLIA, the sensitivity of the established CLIA was 1:128, ASFV-specific serum antibody was identified, and there was no cross-reaction with other swine virus antibodies. After testing 49 clinical serum samples, the consistency rate between the CLIA and the World Organization for Animal Health (OIE) recommended commercial kit was 100%. Thus, this CLIA had a high degree of specificity, sensitivity, and reliability, and could be used as a rapid detection method for epidemiological investigations of ASFV infection.

## Introduction

African Swine Fever (ASF) is an acute, febrile, and highly contagious animal infectious disease of pigs caused by African Swine Fever Virus (ASFV) (1, 2). Its clinical symptoms mainly include skin congestion and cyanosis, severe bleeding of internal organs, high morbidity, high mortality, and a short disease course (2, 3). The virus is transmitted through contact with infected animals and contaminated feed or pollutants, and may also be transmitted by soft ticks; its main hosts are warthog and bushpigs (4). The ASFV belongs to the ASFV family and is a double-stranded linear DNA virus with a tetrahedral structure, a full-length genome of 170~190 kb, 151 open reading frames, and an envelope (5, 6). In addition, there is a variable region at each end of the ASFV genome; its genome has 24 genotypes, which encodes 150–200 proteins (7, 8). The ASFV p54 protein, encoded by the E183L gene, is an important structural protein and is located in the inner envelope of the virus particle; its molecular weight is approximately 29 kDa (9). The ASFV p54 protein plays an important role in the process of viral infection, especially in the conversion of viral proteins into viral envelope precursors through the endoplasmic reticulum membrane (10). The ASFV p54 protein is a late viral protein and thus, transcription of the ASFV p54 gene occurs in the late stage of virus infection (6). P54 is strongly conserved and is a good immunogen, suggesting it may be used as a detection antigen of an ASFV antibody (11).

The chemiluminescence immunoassay (CLIA) was proposed by Halman et al. in 1977 (12). CLIA, which combines a highly sensitive chemiluminescence reaction system with a highly specific immune response system, is currently the most widely used analytical technique (13). The basic principle of CLIA is labeling antigens or antibodies with tracers, such as luminescent agents or catalysts of a chemiluminescent reaction (14). The labeled antigens or antibodies react with the detected substances in a series of immune reactions, and the detected substances are quantified by measuring the luminescence intensity (15). CLIA is divided into three types: chemiluminescence enzyme immunoassay (CLEIA), chemiluminescence labeling immunoassay (CLL), and electrochemiluminescence immunoassay (ECL) (16). CLIA has a high sensitivity, wide detection range, fast reaction speed, short analysis time, and simple operation (16, 17). CLIA is a combination of the chemiluminescence and immunoassay methods(14). With its high sensitivity and specificity, CLIA has continuously contributed to a new understanding of scientific technology and research fields (18). With the rapid development of CLIA in recent years, CLIA has been widely used in the fields of life science research, drug analysis, medical clinical research, food analysis and detection, and animal medicine and environmental management; many important research results have been achieved using CLIA (19). For example, Liu et al. have established a CLIA method for proteins based on recombinant non-structural epitopes that accurately distinguishes cattle infected and inoculated with foot-and-mouth disease virus with high sensitivity (20). D. Chen et al. have compared CLIA, enzyme-linked immunosorbent assay (ELISA), and the passive agglutination method by detecting *Mycoplasma pneumoniae* infection(21). They find that CLIA and ELISA have higher sensitivity than that of the passive agglutination method, and the CLIA results are highly consistent with the ELISA results (21).

Although it has been indicated that CLIA may be used for disease detection with a high sensitivity and specificity, CLIA has relatively few clinical applications in veterinary medicine. In this study CLIA was used to detect ASFV, specifically to establish an assay which was highly sensitive, so as to facilitate clinical detection of ASFV.

## Materials and Methods

### Material sources

ASFV was preserved in the laboratory. Positive sera against pathogenic pseudorabies virus (PRV), Porcine parvovirus (PPV), Porcine reproductive and respiratory syndrome virus (PRRSV), Japanese encephalitis (JE), and classical swine fever virus (CSFV) was prepared in our laboratory. ASFV standard immunized sera (n = 1) and infected sera (n = 33) were obtained from the Laboratory of Animal Health and Epidemiology Center in China. The ASFV ELISA antibody detection kit was purchased from INgezim (Spain). Freund’s adjuvant and goat anti-mouse lgG were purchased from Sigma-Aldrich (Missouri, USA).

### Production of the ASFV p54 protein

The ASFV p54 gene sequence from the Benin 97/1 strain (GenBank ID: AM712239.1) was used to prepare the p54 recombinant protein fragment. The corresponding nucleotide sequence was codon-optimized and synthesized using integrated DNA technology (Wuhan Aoke Dingsheng Biotechnology Co., Ltd. Wuhan, China) to obtain the recombinant plasmid pET-32a (+)-p54. Subsequently, His-tagged truncated p54 constructs were cloned into the prokaryotic expression vector pET-42b (+) to obtain the recombinant plasmid pET-42b(+)-p54. The two plasmids were expressed in BL21(DE3) *Escherichia coli* (*E. coli*) and the AFSV p54 protein was purified using affinity chromatography. The obtained protein was tested by western blot using a HIS-tag, glutathione S-transferase (GST)-tag, and standard positive serum to verify its immunogenicity.

### Monoclonal antibody (mAb) production

A mixture of 100–200 μg ASFV p54-HIS protein and an equal volume of adjuvant was used to immunize 6–8 weeks old BALB/C mice. Mice were immunized three times on the back, with an interval of 2 weeks. Before cell fusion, mice were immunized with 200 μg ASFV p54-HIS protein in the abdominal cavity, euthanized 3 days after the final immunization, and then splenocytes were collected and fused with SP2/0 myeloma cells. Cells were cultured in hypoxanthine-aminopterin-thymidine selection medium (Sigma-Aldrich, Missouri, USA) in 24-well plates after fusion. The recombinant protein ASFV p54-GST was used as the screening antigen, and an ASFV p54-GST indirect ELISA method was established to screen expressed ASFV anti-p54 specific antibodies. Hybridoma cells that produced ASFV anti-p54 specific antibodies were subcloned into single monoclonal cell clones, and the resulting monoclonal cell lines were injected into mice. Prior to this, mice were injected with 500 μL Freund’s incomplete adjuvant. Mouse ascites were collected after 10–15 days and purified using the octanoic acid-ammonium sulfate method to obtain specific ASFV anti-p54 mAbs.

### Western blot analysis

ASFV-infected porcine alveolar macrophage (PAM) cells were harvested at 72 hours post-infection using IP lysis buffer containing protease inhibitors. Cell debris was removed by centrifugation at 15,000 × *g* for 10 min at 4 °C. The cell lysate was mixed with sample buffer (5×) and boiled at 100 °C for 10 min. At the same time, PAM cells not infected with ASFV were harvested as the control. After separation by sodium dodecyl sulfate-polyacrylamide gel electrophoresis (SDS-PAGE), the protein was transferred to a nitrocellulose membrane. The membrane was blocked with 1% bovine serum albumin (BSA) + 1× phosphate-buffered saline (PBS), containing 0.05% TWEEN 20 (PBST) for 2 h at 37 ℃, and incubated with anti-p54 mAb for 1 h at room temperature. After washing with PBST 5 times, the membrane was further incubated with goat anti-mouse IgG secondary antibody for 1 h at room temperature. After washing with PBST five times, the blots were imaged using a multimode reader (Tecan, Austria)

### Immunofluorescence assay (IFA)

The IFA test was performed on PAM cells infected with the ASFV. The cell monolayer was fixed with 4% paraformaldehyde for 15 min after 72 h of infection. Cells were permeabilized by incubating with 0.1% Triton X-100 for 15 min, then blocked with 1% BSA for 1 h, incubated with anti-p54 mAb for 1 h, and then incubated with Alexa-Fluor 594 conjugated goat anti-mouse IgG secondary antibody for 1 h. The nucleus was counterstained with 4’,6-diamino-2-phenylindole, and the fluorescence intensity was observed using a laser confocal microscope.

### Development of the CLIA

A CLIA was established to detect antibodies against ASFV using ASFV p54-HIS and the ASFV anti-p54 mAb obtained in this study. First, plates were coated with ASFV protein p54-HIS. After blocking with blocking solution, ASFV standard serum was added and incubated for 1 h, and then the plate was washed five times with PBST. Next, ASFV anti-p54-horseradish peroxidase mAb was added and incubated for 1 h. After washing with PBST again, Luminol developing solution was added, and the plate was placed in the microplate reader to read the luminescence value. The CLIA cut-off value was evaluated by performing receiver-operating characteristic interactive dot diagram analysis using the SPSS statistical software. CLIA conditions, such as ASFV p54 coating and blocking conditions, sample and conjugate dilution, buffer, and incubation time, were optimized for antibody detection in serum.

### Sensitivity and specificity assay

The standard ASFV positive serum was twofold diluted in sample diluent, and then the CLIA built in this study and a commercial kit were simultaneously used to detect the diluted serum, and the sensitivity of the two methods was compared. Additionally, , PR-, PP-, PRRS-, JE- and CSF-infected sera were also used to evaluate the specificity of the CLIA built in this study and the commercial kit.

### Compliance analysis

The CLIA and the commercial kit were used to test 49 clinical samples simultaneously to evaluate the accuracy of the CLIA established in this study.

## Results

### Generation of mAbs against ASFV p54

The obtained recombinant proteins p54-HIS and p54-GST were subjected to SDS-PAGE, and the two purified proteins were 37 kDa (Fig. 1a) and 43 kDa (Fig. 1b), respectively. Western blotting was performed using an anti-HIS tag mAb, anti-GST tag mAb and ASFV positive serum. P54-HIS showed specific reaction bands with the anti-HIS tag mAb and ASFV positive serum, but not the anti-GST tag mAb (Fig. 2). In contrast, p54-GST showed specific reaction bands with the anti-GST tag mAb and ASFV positive serum, but not the anti-HIS tag mAb (Fig. 3). These results indicated that both p54-HIS and p54-GST had biological activity, and there was no tag cross-reaction. Then, according to the classic procedure of mAb preparation, obtained mouse ascites was purified using the caprylic acid-ammonium sulfate method. As shown in Fig. 4, the purified mAb has a heavy chain at 55 kDa and a light chain at 19 kDa. Finally, a purified anti-p54 mAb was obtained.

**Fig. 1.**
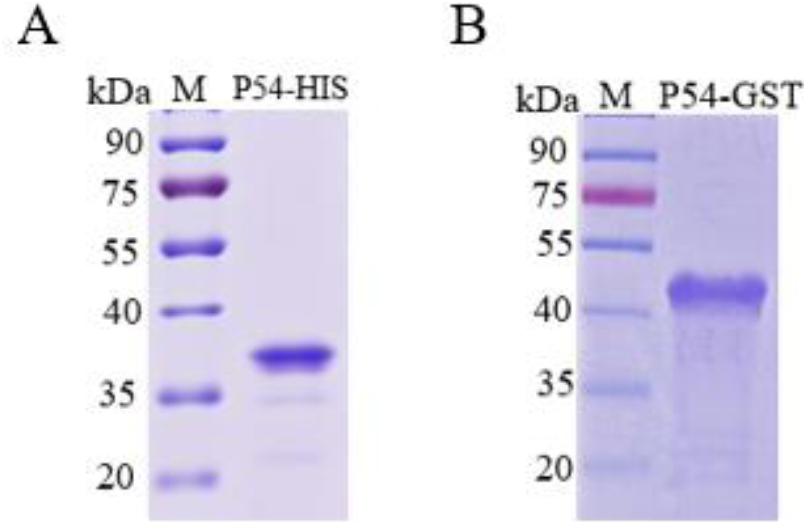
Prokaryotic expression of recombinant p54 proteins. Purified p54-HIS (A) and p54-glutathione S-transferase (GST) (B), as analyzed by sodium dodecyl sulfate-polyacrylamide gel electrophoresis (SDS-PAGE) to identify the expression and purity of recombinant p54 proteins.

**Fig. 2.**
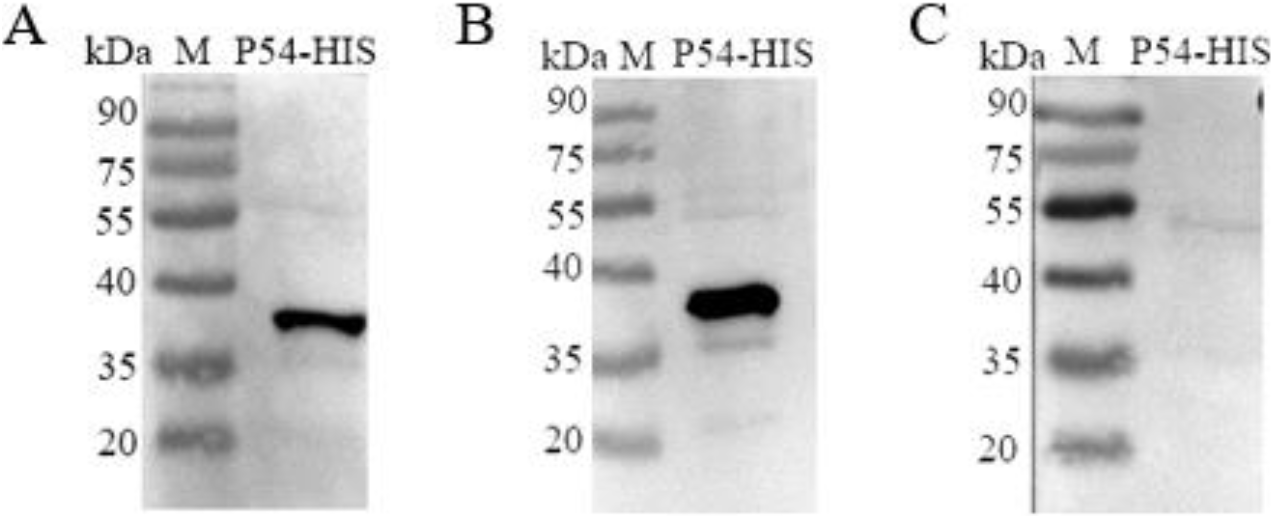
Western blot analysis of recombinant p54-HIS protein. Western blotting has been performed using an anti-HIS tag mAb (A), African swine fever virus (ASFV) positive serum (B), and an anti-glutathione S-transferase (GST) tag mAb (C) to confirm the expression of recombinant p54-HIS protein.

**Fig. 3.**
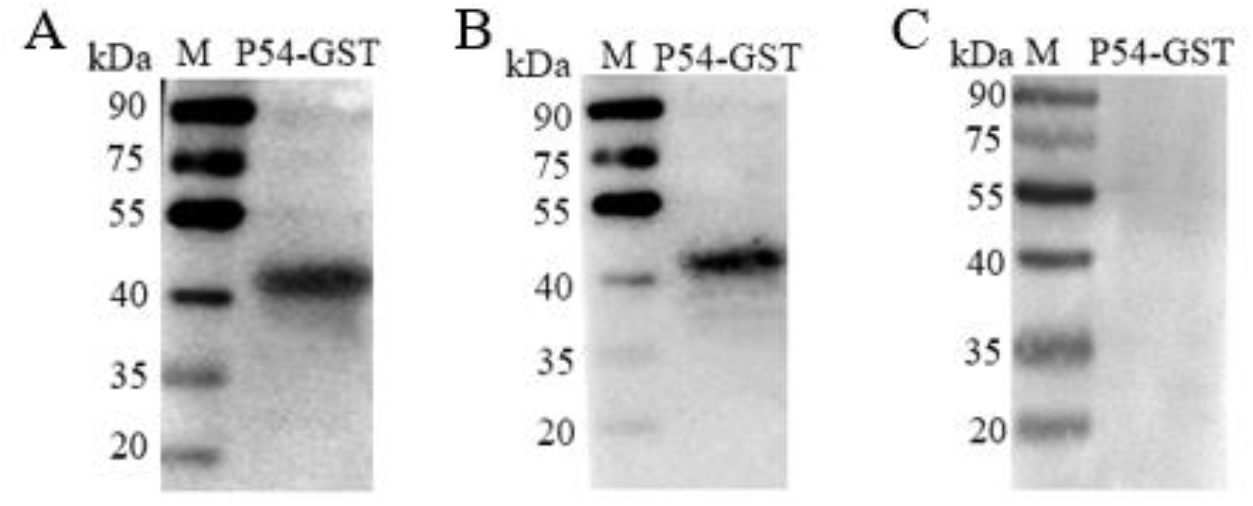
Western blot analysis of recombinant p54-glutathione S-transferase (GST) protein. Western blotting has been performed using an anti-GST tag mAb (A), African swine fever virus (ASFV) positive serum (B), and an anti-HIS tag mAb (C) to confirm the expression of recombinant P54-GST protein.

**Fig. 4.**
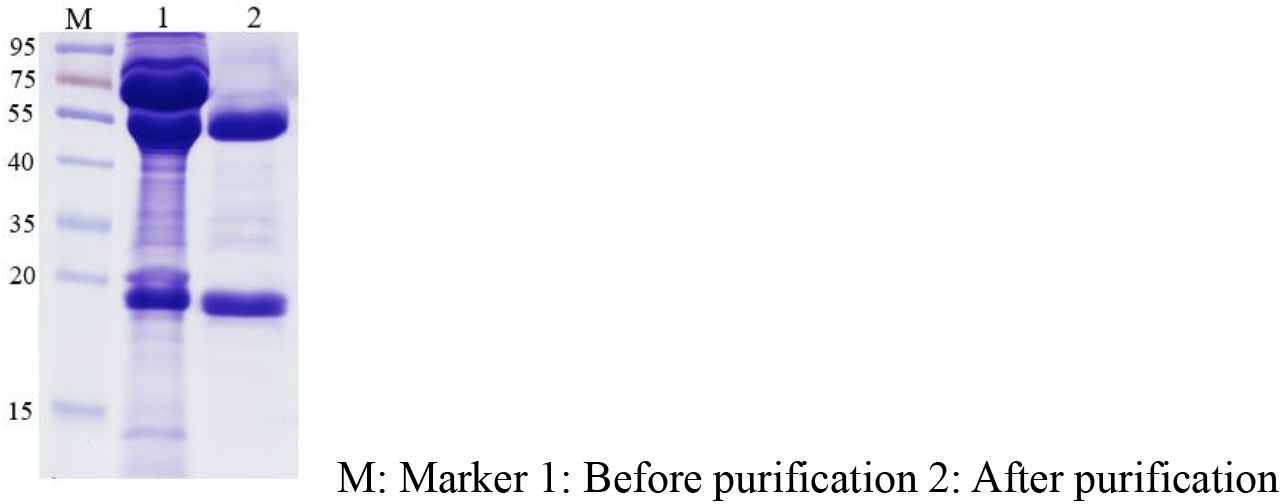
Purification of African swine fever virus (ASFV) monoclonal antibody against protein p54. Harvested ascites are purified using the octanoic acid-ammonium sulfate method process.

### Reactivity of the anti-p54 mAb in ASFV-infected cells

A western blot was performed on PAMs infected with the ASFV. As shown in Fig. 5, PAMs infected with ASFV are detected using the anti-p54 mAb, and there is a specific band at 29 kDa, which is consistent with the size of ASFV p54, suggesting the anti-p54 mAb recognizes ASFV. Further, the reactivity of the mAb in infected cells was validated using an IFA test. Infected cells had a fluorescent signal, indicating the anti-p54 mAb specifically recognized ASFV, while uninfected cells did not have a fluorescent signal (Fig. 6). Thus, the results of western blot analysis and the IFA test demonstrated that the anti-p54 mAb specifically recognized ASFV in infected cells.

**Fig. 5.**
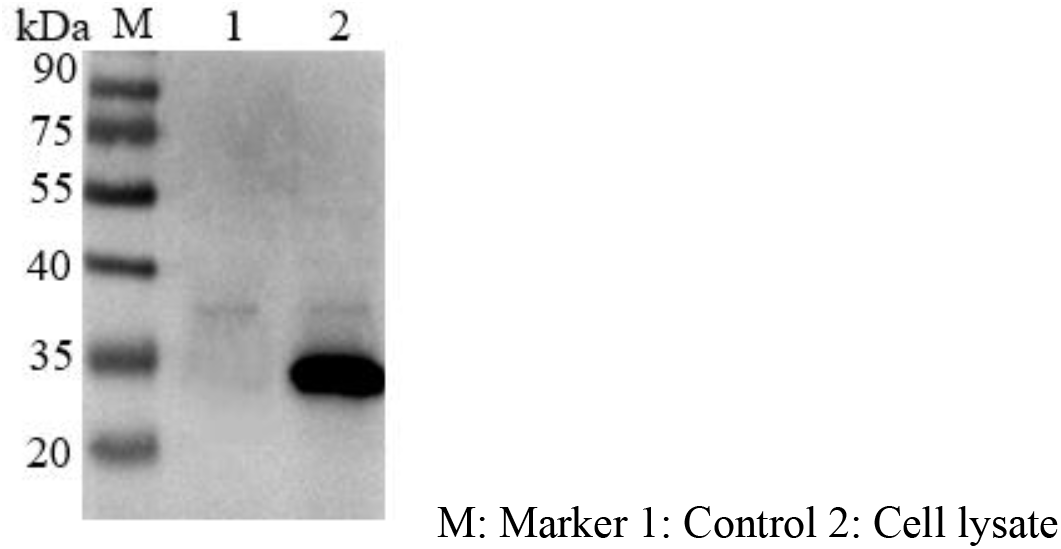
Western blot analysis of monoclonal antibody against protein p54.

**Fig. 6.**
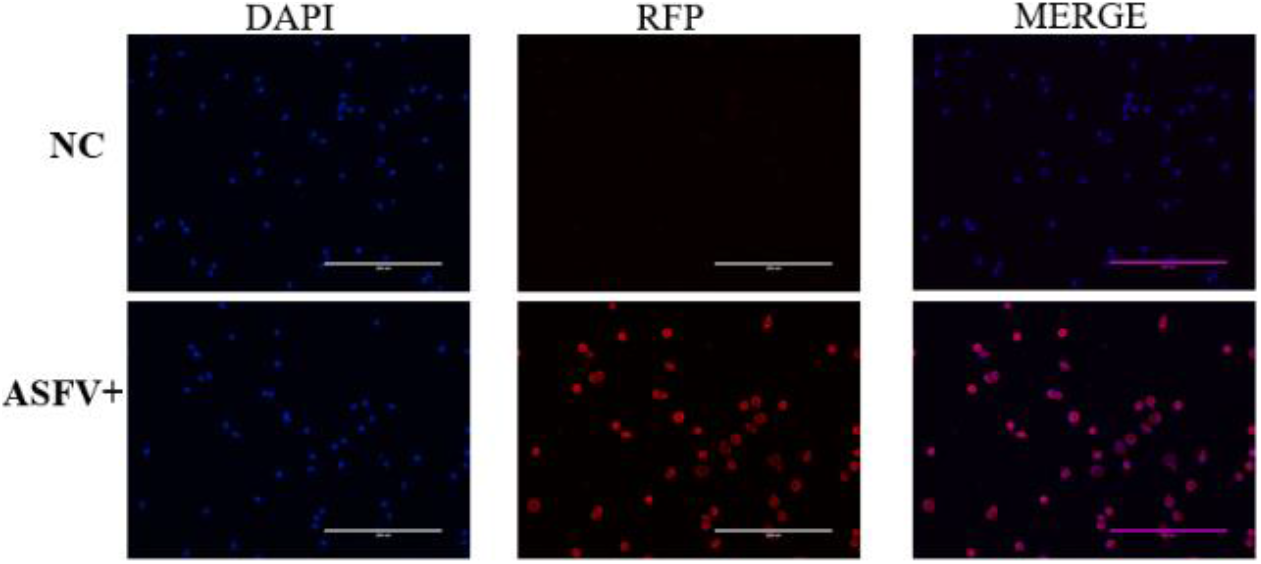
Immunofluorescence assay analysis of anti-p54 monoclonal antibody.

### Optimization of the CLIA protocol

The 96-well microtiter plates were manually coated with 100 μL 0.3 μg/mL p54-HIS in PBS at pH 7.4 and then incubated overnight at 4 ℃, according to the square matrix titration and condition exploration. After incubation, the plates were washed 5 times with PBST, sealed with a solution containing 1% BSA (120 μL/well), and incubated at 37 ℃ for 2 h. Then, serum samples were diluted 1:2, and 100 μL diluted samples were added per well. Positive and negative antibody controls were run in duplicate on each plate. The plates were incubated at 37 ℃ for 30 min and then washed 5 times with PBST. After that, 100 μL peroxidase-conjugated anti-p54 mAb against ASFV was diluted 1:20,000 and added to each well, and the plate was incubated at 37 ℃ for 30 min. The plates were washed again 5 times with PBST, and finally, substrate solution A (Luminol) and solution B (Luminol enhancer; Thermo Fisher Scientific, USA) were added. After 6 min, chemiluminescence values were determined using a chemiluminescence instrument. Medcalc statistical software showed that CLIA’s the percent positive (PP) ritical value was 50%, PP ≥ 50%, sample judged positive; PP ≤ 50%, sample judged negative using the equation PP = (1-CL _sample_ / CL _negative control_) ×100%.

### Comparison of sensitivity and specificity

To evaluate the sensitivity of the CLIA, serum samples with different dilution ratios were tested using the CLIA established in this study and the commercial kit under optimal detection conditions. The sensitivity of the CLIA was 1:128, and the sensitivity of commercial kits was 1:16. Therefore, the sensitivity of CLIA was 8 times that of the commercial kits. (Fig. 7a). Meanwhile, the CLIA specificity was analyzed using specific serum samples containing other common swine viruses, including CSFV, PRRSV, JEV, PRV, and PPV. The test results of CLIA and the commercial kits were both negative and the results of the two were consistent, demonstrating that the CLIA had good specificity (Fig. 7b).

**Fig. 7.**
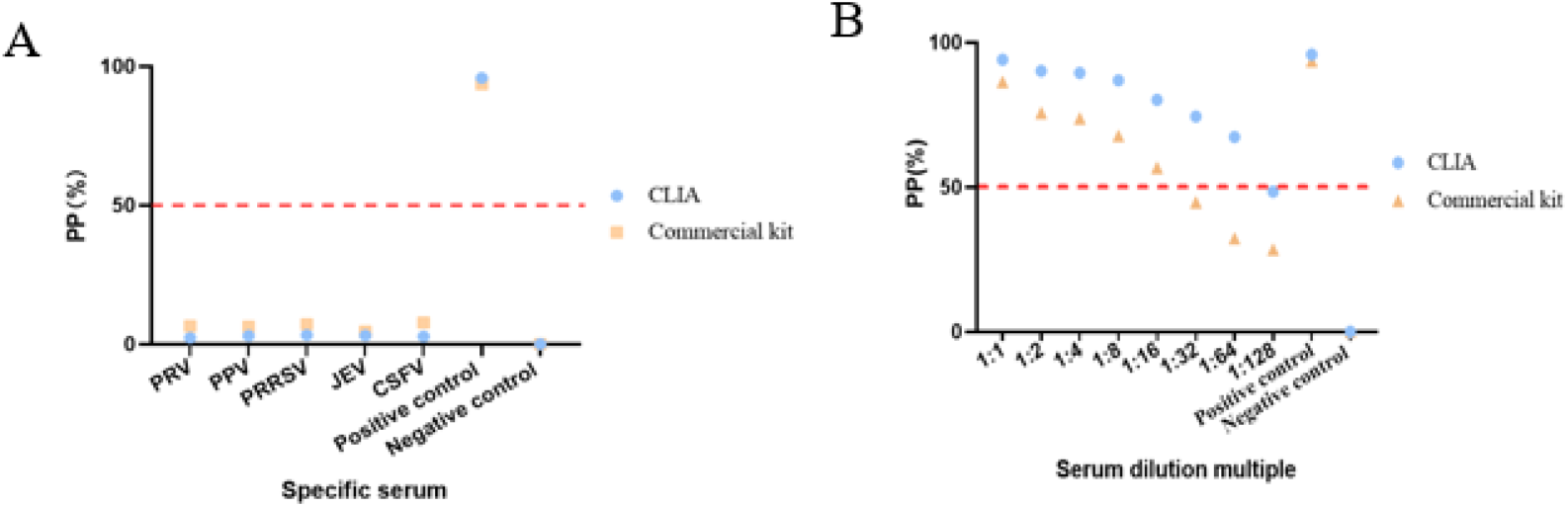
The specificity (A) and sensitivity (B) of the chemiluminescent immunoassay (CLIA) and the commercial kit have been simultaneously tested.

### Comparison of Coincidence test

To evaluate whether the detection method established in this study was suitable for ASFV antibody detection in a clinical application, a coincidence test was performed between the CLIA and commercial kits. As shown in Table 1, the CLIA and commercial kits have been used to test 33 positive samples and 16 negative samples. CLIA test results were 100% consistent with the commercial kits test results. Therefore, CLIA could detect ASFV antibodies.

**Table 1.**
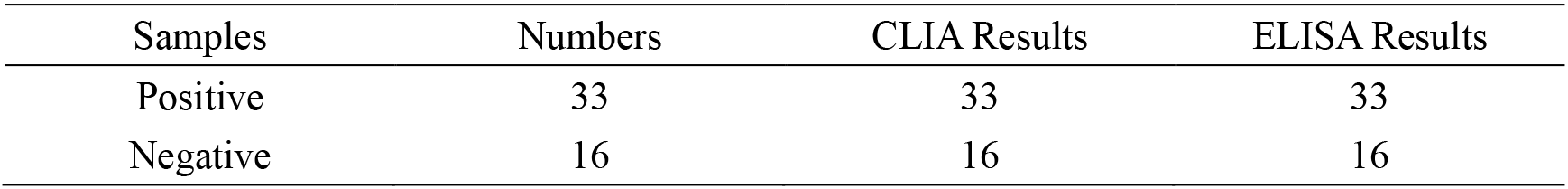
Results of chemiluminescent immunoassay (CLIA) and enzyme-linked immunosorbent assay (ELISA)

| Samples | Numbers | CLIA Results | ELISA Results |
| --- | --- | --- | --- |
| Positive | 33 | 33 | 33 |
| Negative | 16 | 16 | 16 |

## Discussion

ASFV was first reported in Kenya, Africa in 1921, and was introduced to Portugal, a European country, in 1957. From the 1970s to the 1980s, ASFV was identified in many countries in Europe and America (22). China reported African swine plague for the first time in 2018, and then the disease spread rapidly, causing huge economic losses to the pig industry and the national economy (23). ELISA is the gold standard recommended by OIE for detecting antibodies against ASFV, and since the outbreak of ASF in China, indirect ELISAs and competitive ELISAs using various antigenic genes to monitor serum for ASF infection have been widely established (24, 25). Generally, compared to an indirect ELISA, a competitive ELISA has better specificity, but worse sensitivity (26). Previous studies have shown that CLIA is more sensitive and specific than ELISA (16, 19). However, CLIA is rarely used in veterinary clinics, especially for ASF. Therefore, in this study, ASF was combined with CLIA to develop a specific, sensitive, and reliable detection method.

The OIE approved the current ASFV antibody-based test based on the use of serum as the test sample and live virus as the antigen, which requires a level 3 biosafety facility to spread and handle live virus (7). However, CLIA detects ASFV antibodies based on recombinant antigens, which avoids the risk of using live viruses, and achieves the same effect. In addition, when comparing CLIA and commercial ELISA kits in terms of specificity and sensitivity, the results obtained are consistent with previous research conclusions (16), CLIA’s specificity and sensitivity are higher than commercial ELISA detection kits.

There are occasional outbreaks of ASF, and worse, recovered ASFV carrier pigs and persistently infected wild pigs constitute the biggest challenge to controlling the disease (7). However, there remains no effective vaccine to prevent ASF worldwide, which means monitoring is essential. Pigs surviving a natural infection will develop antibodies against ASFV 7–10 days after infection, which are detected for a long time(7). Additionally, serum specimens are easy to collect and have high diagnostic information. The detection of antibodies to ASFV in pig serum suggests that the pig has been infected by ASFV. Therefore, the current research developed a simpler and faster antibody detection method based on effective and low-cost specimen collection.

This method provides convenience for testing convalescent and infected pigs serologically. While qPCR is the gold standard recommended by OIE to detect antigens (27), qPCR occasionally produces false positives due to inappropriate sample handling. Therefore, the CLIA from this study could be combined with qPCR to screen healthy and negative pigs, which would avoid increased losses and be more conducive to the recovery of the pig industry.

In summary, this study developed a CLIA method based on an ASFV anti-p54 mAb, which had high sensitivity, specificity, and reliability. This CLIA method requires a shorter detection time, which is more suitable for the epidemiological investigation of ASFV compared to that of other ELISA methods, may be used in combination with other diagnostic testing methods, including etiology and serology techniques, and may even facilitate the development of vaccines to prevent and control ASF.

## Acknowledgements

This work was supported by The Major Projects of Technological Innovation in Hubei Province (2019ABA089).

